# Phytochromes Enable Social Behavior in Marine Diatoms

**DOI:** 10.1101/2024.09.18.613651

**Authors:** Joan S. Font-Muñoz, Marianne Jaubert, Marc Sourisseau, Idan Tuval, Benjamin Bailleul, Carole Duchêne, Gotzon Basterretxea, Angela Falciatore

## Abstract

The phytochrome superfamily, a group of proteins that enable some organisms to detect changes in light intensity and quality, is widespread in terrestrial and marine microbes, fungi, algae, and plants. In terrestrial plants, these photosensory receptors monitor variations in the light environment by sensing red (*R*) and far-red (*FR*) regions of the spectrum and trigger important developmental, metabolic, and physiological responses. However, the role of these photosensors in marine microbes, living in environments where, due to absorption of water molecules *R* and *FR* radiation does not penetrate beyond the upper few meters, remains controversial. Here, we investigate the role of phytochromes in light perception of the marine diatom *Phaeodactylum tricornutum* and their involvement in light-driven collective behavior. We perform experiments comparing the social conduct of wild-type and phytochrome knock-out strains to different light wavelengths. Our results show that cell movements become synchronized in a coordinated wobbling dance upon activation of their phytochromes by blue or far-red light, therefore, demonstrating the key role of phytochrome in light-mediated diatom collective behaviour. Furthermore, our experiments suggest that the observed phytochrome-mediated concerted dance implies a form of intercell communication, proposedly mediated by variable R/FR autofluorescence emission in the frequency range of diatom wobbling movements. Our findings provide new insights into communication pathways in aquatic microorganisms and emphasize the importance of social conduct in the sea at all ecological levels.

## INTRODUCTION

Photosynthesis is an essential biological process that sustains life on Earth, by enabling autotrophic organisms to transform solar energy into chemical energy, which in turn serves as a vital source of sustenance for other living organisms. The effectiveness of photosynthesis relies, among other factors, on the ability of autotrophs to adapt to fluctuating environmental sunlight conditions. To this end, they have evolved highly specialized sensing molecules that allow the perception of incoming environmental signals and transduction into biochemical outputs that shape organism responses (*1, 2*).

Phytochromes are a group of photoreceptor chromoproteins that enable plants and some photosynthetic and non-photosynthetic microbes to detect and adapt to changes in light intensity and quality (*3*). In terrestrial plants, phytochromes (PHYs) are capable of sensing light in the red (*R)* and far-red (*FR)* regions of the spectrum (*4*). PHYs provide essential plasticity to plant survival, by activating specific signaling cascades and allowing active adjustment of vital processes such as germination, growth, development, phototropism, and related metabolic pathways (*5, 6*). Plant PHYs are also capable of perceiving changes in the *R/FR* ratio of ambient light caused by neighboring vegetation, initiating shade avoidance and acclimation reactions (*7*). In some human pathogenic bacteria, PHYs-mediated light signals also integrate quorum-sensing responses to control collective behaviors and diverse lifestyles (*8*).

PHYs have also been recently found in a broad range of aquatic microorganisms. Their discovery first came as a surprise because *R* and *FR* wavebands derived from sunlight radiation are scarce in aquatic environments, due to their rapid attenuation by water absorption (*9, 10*), opening questions about the mechanisms of action and function of these photoreceptors (*11*). The more recent characterization of phytochromes from diverse eukaryotic algae provided some answers to this conundrum. Extensive spectral tuning and blue-shifted absorption spectrum have been described in PHYs of some prasinophyte and glaucophyte species, leading to the hypothesis of an adaptation of the photoreceptor absorption properties to the blue-rich aquatic environment (*12*). On the contrary, conserved *R* and *FR* light absorption spectra have been described for the phytochrome of diatoms (*13*), one of the most prominent and diversified phytoplankton groups in the ocean which are distributed worldwide and play a key role in global biogeochemical cycles (*14, 15*). Unforeseen roles for *R* and *FR* PHYs-mediated detection have been hypothesized, such as the perception of fluorescent light emitted by nearby photosynthetic congeners, potentially providing valuable information on cell densities, for example, during algal blooms (*13, 16, 17*), and stimulating various aspects of their cellular metabolism, such as iron uptake (*17*). Recent studies, however, indicate that the Diatom Phytochromes (DPH), despite the conserved *R* and *FR* light absorption (*13*), can also exhibit photoreversible responses across the whole available underwater light spectra, suggesting an additional potential role of DPH as an optical depth sensor (*18*) but not ruling out other functional roles.

Diatom lifestyle diversity and survival success in contemporary oceans are attributed to their exceptional ability to adapt to highly dynamic aquatic environments using efficient mechanisms to respond to environmental changes (*19*–*22*). For example, several diatom species can cope with highly variable light conditions, suggesting that diatoms are capable of perceiving, responding, and likely, anticipating light variations and that they possess suited underlying molecular systems mediating light responses (*23, 24*). Recent studies have also shown that several pennate marine diatoms are indeed capable of perceiving and using external light signals to synchronize intracellular processes driving their unsteady sinking behavior (*25, 26*). However, the molecular mechanisms underpinning these photoresponses and the role of PHYs in light-mediated behavior at the population level are not yet established.

Here, we use several independent *P. tricornutum dph* knock-out lines, together with laser diffractometry, to evaluate the role of *photosensing* in the light-mediated development of collective cell behavior (*27*–*30*). We experimentally demonstrate that photoreception by DPH is required to establish a coordinated social pattern in populations of *P. tricornutum* suspended in the water column. The possibility that endogenous light signals are used by microbial organisms to perceive and communicate with neighboring homologous cells to develop and propagate coordination is also discussed.

## RESULTS AND DISCUSSION

Pelagic diatoms with elongated shapes characteristically display coherent vertical cell orientation and synchronized wobbling when sinking under continuous illumination (*25, 26, 31*). In the case of nearly elongated cells, this behavior can be characterized by laser diffractometry (LD) using the relative variation of two size bands associated with the major and minor cell axis lengths (*32, 33*). Due to the 2D projection of the LD measurements, the *Ratio* between the signals of these two size bands is a scalar proxy for the mean orientation of the cell population (see, M&M). As shown in Figure 1 (see also Fig. S1), in the diatom *P. tricornutum*, the *Ratio* time series displays a pronounced wavelength dependence of synchronized wobbling, with strong responses in the *B* and *FR* regions, but minimal responses elsewhere in the light spectrum. This spectral dependence is consistent with that described by Duchêne et al. (*18*) for DPH-mediated gene expression (Fig. 1B). No coherent wobbling is observed in *dph-knock-out* strains, while their respective control lines (*Tc*, transformed but non-mutated in *DPH*; (*13*)) exhibit a synchronized collective response under *B* light, which is indistinguishable from that observed in the wild-type (*WT*; Fig. S2). These results strongly support that DPH photoperception is strictly required for light-induced synchronized cell wobbling.

**Figure 1.**
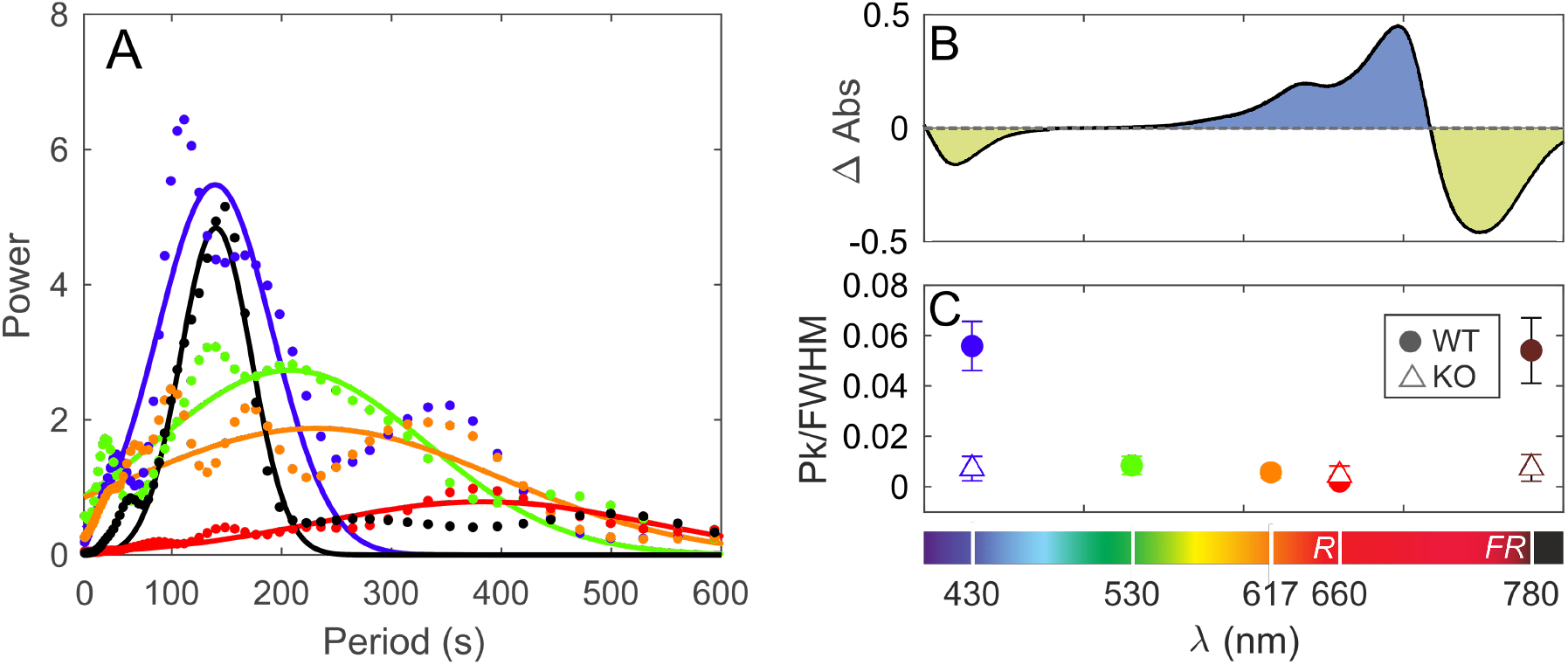
Continuous light experiments. A) Power spectrum of *P. tricornutum* (WT) *Ratio* obtained with laser diffractometry experiments (see Fig. S1 for the original time series). Colors indicate the wavelength with which cells were illuminated (430, 530, 617, 660, and 780nm). Light irradiance was kept constant for all wavelengths (8 *μ*mol photons m^-2^ s^-1^). B) DPH deactivation/activation spectrum (*Pfr-Pr*) resulting from absorption differences between the synthesized active state *Pr* of DPH and the inactive conformation *Pfr*. C) Peak amplitude of the power spectrum (shown in A) normalized by the width of the power spectrum for the wavelength of the lighting used. Solid circles correspond to experiments with wild-type (WT) and transformed but *DPH* non-mutated strains and empty triangles correspond to experiments carried out with *DPH*-knockout strains (KO).

A key feature of PHYs-mediated control is its ability to scale the magnitude of a response according to wavelength ratio. Since, in *P. tricornutum*, DPH was shown to be activated by *B* or *FR* and inactivated by *R* light (*18*), we tested whether the described collective behavior is hindered by changing the relative intensity of *B* and *R* light. Starting with the intensity of *B* light used in Fig. 1, we increasingly intensified *R* light and observed inhibition of the synchronized wobbling amplitude (Fig. 2, see also Fig. S3) while wobbling frequency was unaffected (Fig. 2A). To confirm that this was a response to *R* light and not just the consequence of modified light quantity, we also performed control experiments with increased *B* light intensity and observed no change in the response (Fig. 2B). Our results suggest that, similarly to the proposed *in situ* modulation of DPH activation by optical depth throughout the water column (18), synchronized wobbling is also modulated by blue-to-red ratio. As a consequence, it would be favored at depth, an environment dominated by *B* light.

**Figure 2.**
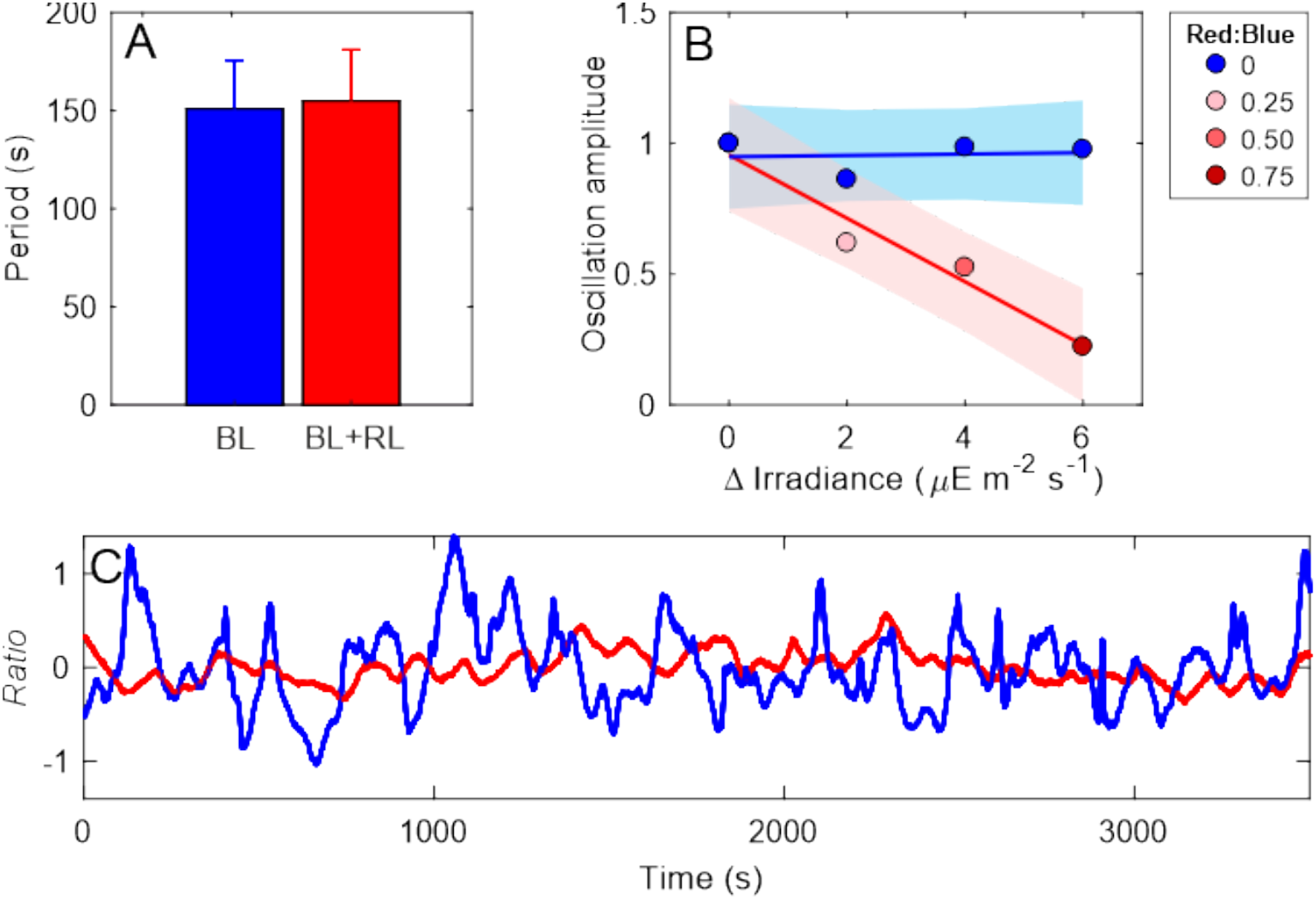
Diatom response to composed continuous blue and red light. A) Mean period of oscillation of WT cells for experiments with different ratios of red:blue light and the *B light* controls. B) Normalized oscillation amplitude for experiments with enhanced light (under *B light* background of 8 *μ*mol photons m^-2^ s^-1^) at different ratios of red:blue light and the controls with *B light*, color bands indicate the 95% confidence band. C) Normalized *Ratio* time series for experiments with 0.75 red:blue ratio and *B light* control of the same intensity.

While the triggering mechanisms of the described cell behavior rely on ambient *B* light availability and detection by DPH, the mechanism through which independent cells freely suspended in the water column coordinate their movements remains challenging. Font-Muñoz et al. (25) framed the question within the context of the theory of weakly coupled oscillators for phase locking (*34*) which requires a harmonic coupling signal compatible with the cell wobbling frequency (ω; Fig 3). It can be speculated that either hydrodynamic physical disturbances in the flow field surrounding individual cells or the modulation of unknown dissolved infochemicals govern these cell-to-cell interactions (*35*–*37*). However, these are unlikely to generate coherent forcing at the involved spatial and temporal scales. Alternatively, *B* light excited cell autofluorescence provides a plausible explanation for intercellular communication inducing the observed synchronized movements. Autofluorescence emission by photosystem II is a by-product of photosynthesis that is cost-efficient and almost instantaneous, and depending on cell morphology, can provide a cell density dependent but directional signal of the correct frequency (*16, 17*). Although in the sea the intensity of fluorescence emission is always very small compared to the ambient light, it could still convey precious information and be perceivable locally by neighbor cells as a weak coupling signal above the *R* light background. Indeed, the measured emitted chlorophyll fluorescence signal in each arbitrary direction is modulated by the cell’s relative orientation, thus exhibiting a time modulation at the exact frequency of the cell wobbling oscillation, as displayed in Figure 3. This periodic light signal comprises both *R* and *FR* spectral components, whose temporal signals are highly similar (Fig. 3B). It is worth noting that while *dph-*knockout strains autofluoresce with the same spectral qualities as *WT* -at room temperature-, they do not exhibit a population level coherent fluorescence signal modulation. It should also be stressed that illuminating an algal suspension with an exogenous monochromatic light does not guarantee a homologous light environment since both Raman scattering and, more intensely, chlorophyll autofluorescence cause the emission of photons shifted in frequency relative to the incident light, thereby modifying the spectral composition of the surrounding light field (*16, 38*). Exogenous light and chlorophyll-emitted light signals cannot easily be decoupled in experiments using dense microalgal suspensions. This opens the question of whether DPH are mediating cellular responses only to exogenous ambient light or whether they are also involved in the perception of time modulated autofluorescence.

**Figure 3.**
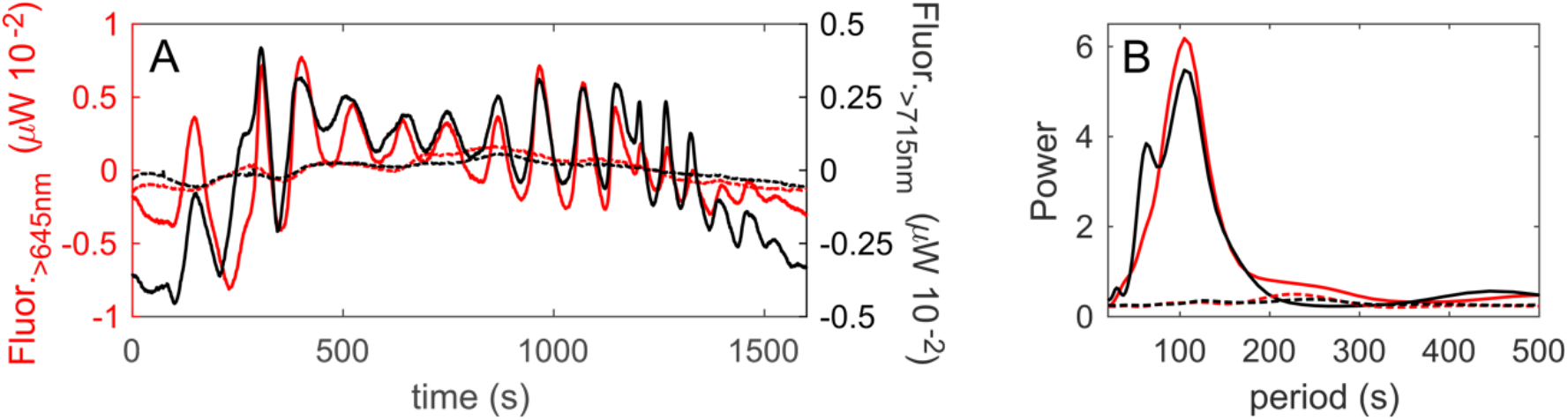
Natural fluorescence emission. A) time evolution of the fluorescence emitted at two different spectral bands (λ>645 and λ>715 nm) in a suspension of *P. tricornutum* (*WT* solid lines, *DPH*-knockout dashed lines) illuminated with 430nm light (1mW) and (B) the corresponding power spectrum.

The idea that PHYs enable algae to sense neighboring cells through the perception of cell autofluorescence is not novel (*13, 16*) but, to the best of our knowledge, has yet to be confirmed. We set out to test this hypothesis by interrogating cells only with pulsed *R* and *FR* light that mimic the measured fluorescence signal naturally emitted by *P. tricornutum* cells (see, Fig. 3A). In Figure 4 we show a striking contrast in the response of the *WT* and the *dph-*knockout strains to the same pulsed stimulus, both under *R* and *FR* illumination. While *WT* and *Tc* cells are readily entrained by pulsed light, the wobbling of *dph-* knockout strains remains largely incoherent (Fig. 4A and B). These results indicate that DPH are not only essential for triggering collective behavior under constant light illumination. They are also key to the synchronization process, potentially implying their involvement in the ability of cells to perceive the light emission of their neighboring congeners and hereby contribute to the observed collective behavior by propagating the signals in the population. Full characterization of this function and its ecological relevance remains to be elucidated.

**Figure 4.**
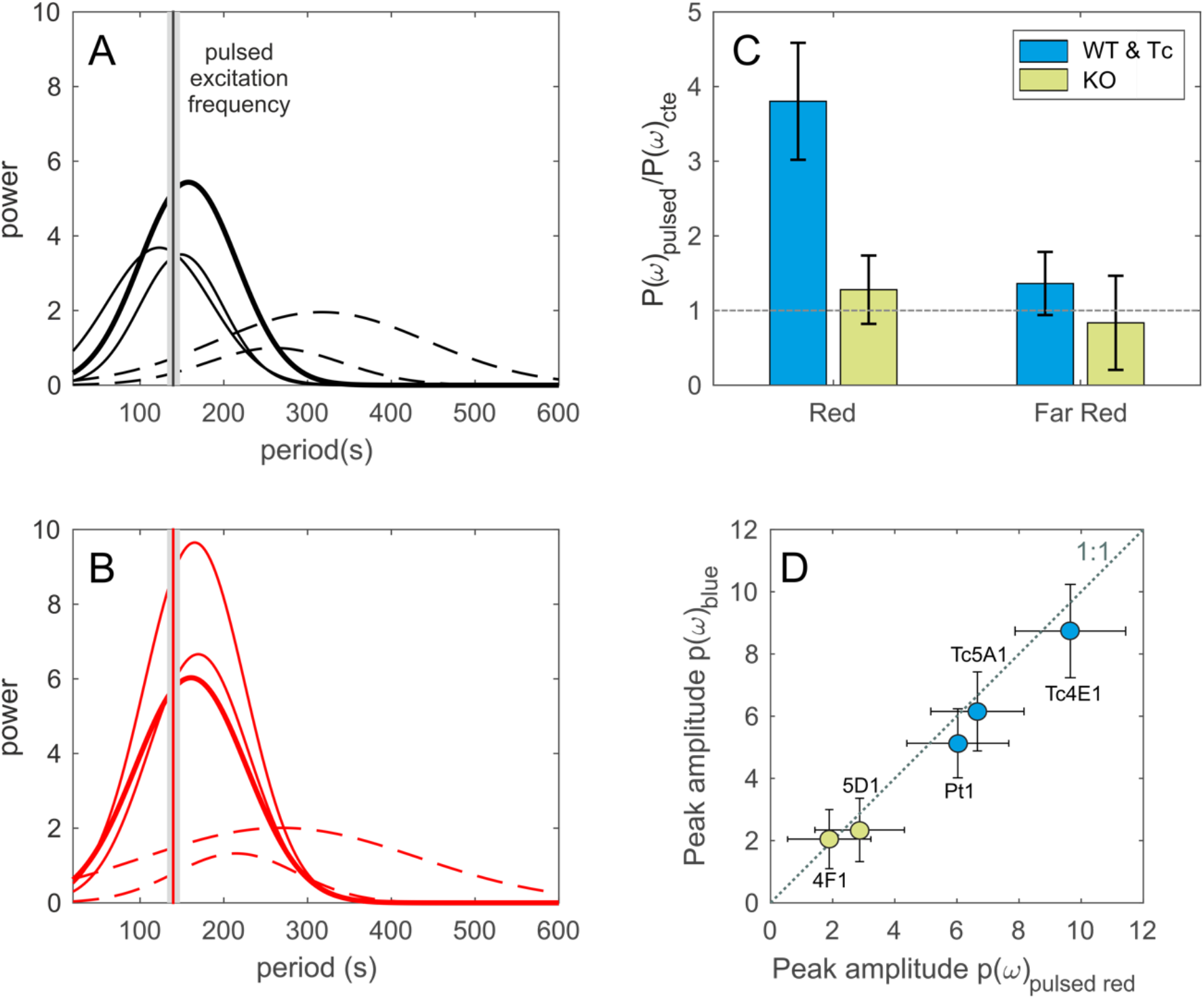
Pulsed light experiments. Power spectrum of *Ratio* for experiments that mimic natural fluorescence emission. The thick solid lines correspond to the *WT* strains, the thin solid lines correspond to the *Tc* strains, and the dashed lines correspond to the *DPH*-knockout strains (*KO*). The grey band indicates the period of light oscillation. Panel A shows experiments with far-red light (780nm), and panel B shows experiments with red light (660nm). C) Differences in collective behavior when using pulsed light versus continuous light: comparison of the peak amplitude of the power spectrum of *Ratio* under pulsed light (red or far-red, ∼140s) to the peak amplitude under constant light (red or far-red), values close to 1 indicate similar collective behavior. D) Peak amplitude of the power spectrum of experiments with constant blue light compared to pulsed red light (∼140s).

Finally, it is important to note that, while a collective photoresponse is not elicited by illumination under constant *R* light in our experiments (see Fig. 1C), a periodic modulation of the signal with frequencies around natural wobbling frequencies (∼140s) suffices to induce strongly coherent cell responses (Fig. 4C). In fact, these are as coherent as those observed under constant *B* (or *FR*) illumination (Fig. 4D and Fig. S4B). Notably, *DPH-knockout* strains do not exhibit collective responses in either case. This differential photoresponse, not observed at any other wavelength, hints at a possible unknown DPH signal transduction pathway, and PHYs-facilitated mechanism, for light perception under quickly fluctuating light environments.

## CONCLUSION

We have found that phytochromes are involved in light-driven coordinated diatom wobbling, a collective behavior modulated by the effective wavebands activating DPH (*18*), and therefore enhanced in *B* light enriched environments. Although the function of this collective dynamics in the dilute marine environment is still unclear, we can speculate it could influence key ecological processes such as sedimentation rates, light harvesting, or increased cell-cell contacts, thereby favoring sexual reproduction in pelagic environments (*25, 26*) especially in the deep, non-turbulent layers of the water column (*31, 39, 40*).

The analysis presented here immediately suggests a number of specific experimental investigations. Chief among them are detailed studies of DPH involvement in the photoresponse to pulsed R light. This can be tested with experiments at the single cell level in which photoresponses to exogenous light can be readily distinguished from signaling from neighboring cells. Open issues include the origins of this regulation, the comprehensive characterization of the perceived stimuli, along with the possible interplay of these processes with other biotic and abiotic signals, as those observed in the quorum sensing of bacteria due to high cell densities (*41*). Additionally, our results under pulsed *R* and *FR* light suggest that oscillations of autofluorescence by cell wobbling could be involved in the propagation of the *DPH*-mediated response. Nonetheless, earlier observations of collective oscillations in pennate diatom species that are not known to express DPHs (25), suggest that DPH may function in coordination with other, yet undiscovered photoreceptors. Overall, our findings provide strong evidence of the relevance of light communication in shaping microorganism interactions and social behavior, and, at a higher hierarchical level, as a structuring element of life in the ocean.

## Acknowledgments

This research was supported by the ISblue project, Interdisciplinary graduate school for the Blue Planet (ANR-17-EURE-0015), and co-funded by a grant from the French government under the program “Investissements d’Avenir’’ and a grant from the Regional Council of Brittany (SAD program). J.S.F-M was supported by funding from “Margalida Comas” postdoctoral fellowship PD/018/2020 from Govern de les Illes Balears and Fondo Social Europeo. Funding from PID2022-143018NB-I00 and PID2019-104232GB-I00 grants from the Spanish Ministerio de Ciencia e Innovación (MICINN), the Agencia Estatal de Investigación (AEI), and the H2020-MSCA-ITN-2020 PHYMOT is acknowledged. The present research was carried out within the framework of the activities of the Spanish Government through the “María de Maeztu Centre of Excellence’’ accreditation to IMEDEA (CSIC-UIB) (CEX2021-001198-M). MJ acknowledges the CNRS MITI interdisciplinary program ‘Lumiere et Vie’ and ‘80PRIME’. BB acknowledges support from the European Research Council (ERC) PhotoPHYTOMIX project (grant agreement No. 715579) and ERC Proof-of-Concept PALMADS (grant agreement No. 101158298). AF was supported by funding from the Fondation Bettencourt-Schueller (Coups d’élan pour la recherche francaise-2018), the “Initiative d’Excellence” program (Grant “DYNAMO,” ANR-11-LABX-0011-01).

## Supplementary Materials

### MATERIALS AND METHODS

#### Cell cultures

The wild-type *Phaeodactylum tricornutum* cells and transgenic lines were grown in f/2 medium (*42*) at 18°C under a 12-h-light/12-h-dark cycle and 80 *μ*m^-2^ s^-1^, in algal incubators. The cultures were transferred once a week (dilution factor x8) to keep the cells healthy and in their exponential growth phase at maximum concentrations. All the experiments were done at concentration of 3×10^8^ cells/L, with cells in exponential phase of growth. *DPH* knockout lines were obtained by TALEN-mediated genome editing as described in Fortunato et al. (*13*). Their respective *DPH* non-mutated lines (derived from the same cell transformed with TALEN vectors but not having undergone mutagenic event on *DPH* gene) were used as control for *DPH* specific phenotype (*13*).

#### Experimental set-up

Experiments were carried out in an in-house built set-up for the concomitant measurement of particle size distribution, orientation and chlorophyll autofluorescence. The small volume (∼100ml) flow through chamber from a LISST-100x laser diffractometer (Sequoia Scientific) was modified to incorporate an extra modulated (spectrally, via a monochromator, and temporarily) light-line, orthogonal to the laser diffraction axis coupled to two power-meters (PM16-130 from Thorlabs). This configuration allows for the interrogation of a wide range of experimental conditions and the simultaneous measurements of sub-second variations in cell orientation and light emission (*25*).

Before each experiment, the LISST-100X chamber was pre-cleaned and filled with the desired v/v concentration of cell culture. Agitation was induced by a 2 cm long magnetic bar powered by the built-in speed controller of the chamber at minimum speed. Experiments consisted in an initial mixing phase where agitation was turned on (100 s) followed by a sedimentation phase where the system could evolve without any external disturbance.

*Diatom response to light:* to study the effect of light conditions on cell orientations the cells and collective behavior were illuminated with continuous light of different wavelengths (430, 530, 617, 660 and 780 nm, mod. M430L5, M530L4, M617L5, M660L4 and M780L3 from Thorlabs) with an intensity of 8 *μ*E m^-2^ s^-1^ for 2h.

*Diatom response to composed blue and red light:* To study the cells responses of cells to composed continuous red and blue light, cells were illuminated with 430 nm at 8 *μ*E m^-2^ s^-1^ and 660 nm at different intensities (2, 4 and 6 *μ*E m^-2^ s^-1^). We performed experiments using monochromatic blue light at the same total intensities (10, 12 and 14 *μ*E m^-2^ s^-1^) as control. We used a Longpass dichroic mirror 550 nm Cut-On (DMLP550 from Thorlabs) to combine the two light sources.

*Diatom response to pulsed red and far-red light:* to study the response of *P*.*tricornutum* to pulsed red light, cells were illuminated by a 660 nm and 780 nm LED (M660L4-C and M780L3, Thorlabs) through a neutral density filter (ND10A Thorlabs) used to attenuate light intensity down to a maximum value of ∼8 *μ*E m^-2^ s^-1^. This intensity was then modulated sinusoidally with periods of cell wobbling natural frequency (∼140 s).

*Fluorescence signal variations measurements:* variations in the cells’ chlorophyll autofluorescence were measured by illuminating the culture with an excitation blue light (430 nm at 1mW) for 45min while the cells settled without disturbance. The emitted *R* and *FR* light coming from the cells was measured every second by the orthogonal power-meters (PM16-130 from Thorlabs) via a high pass *R* and *FR* filters (FGL645M, Thorlabs; λ greater than 645nm and FGL715M, Thorlabs; λ greater than 715nm).

#### Laser diffractometry

Laser diffractometry (LD) measurements were used to obtain the particle volume concentration by size ranges (i.e., volume of particles in the seawater per unit volume of seawater) using a technique based on laser diffraction theory. The LISST-100X employed in this study uses a 670 nm collimated laser beam to illuminate the suspended particles and a 32-ring detector measures the intensity of the scattered light, corresponding to 32 different size classes logarithmically spaced. The angular pattern of optical scattering depends on the physical characteristics of the particles (e.g., size, shape and orientation), and is used to calculate the particle volume concentration in these size classes (*43*). LD can be used to adequately characterize the different dimensions of non-spherical particles in specific orientations (*28, 29, 31*). Using this property, a method to infer the preferential orientation of particles in a suspension has been recently described (*30*). Hereby, we use this method to characterize the preferential orientation of diatoms under distinct lighting conditions: LD measurements of nearly spheroidal cells are interpreted in terms of the relative variation of two size bands representing the known major and minor cell axes (r_1_ and r_2_) of *P. tricornutum*. As cells orient in different directions, the signal in the size bands representing these axes presents an opposite behavior. Hence, we can confidently compute the ratio between the signal from each of these two size bands, Ratio(t)=(∑VD_1_)/(∑VD_2_), as a scalar proxy for cell orientation (*25, 30, 31*).

**Figure S1.**
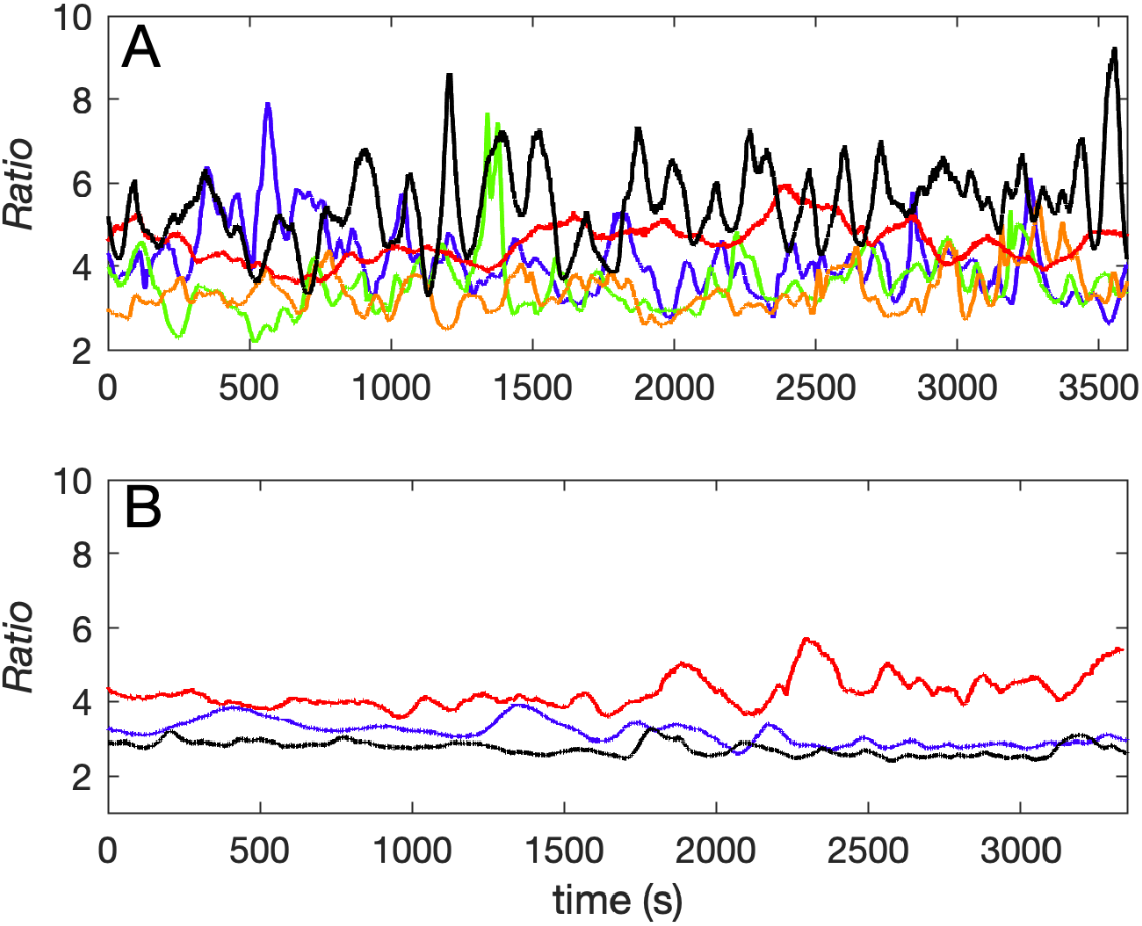
Time series of *Ratio* obtained in *P. tricornutum* experiments performed with A) wild-type strains and B) *dph*-knockout lines. Colors indicate the wavelength with which cells were illuminated (430, 530, 617, 660, and 780 nm).

**Figure S2.**
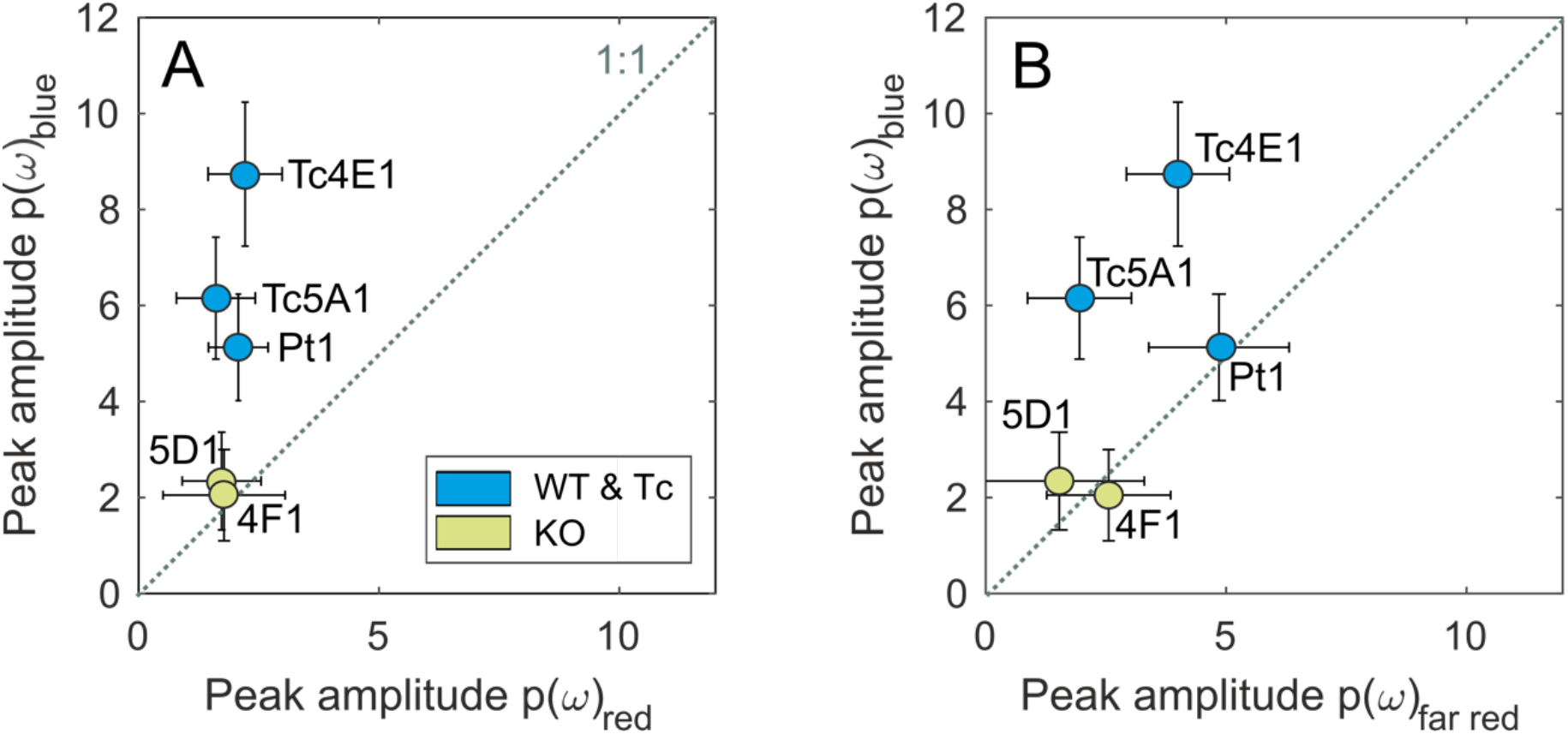
A) Peak amplitude of the power spectrum for experiments with constant blue light compared to constant red light. B) Peak amplitude of the power spectrum for experiments with constant blue light compared to constant far-red light.

**Figure S3.**
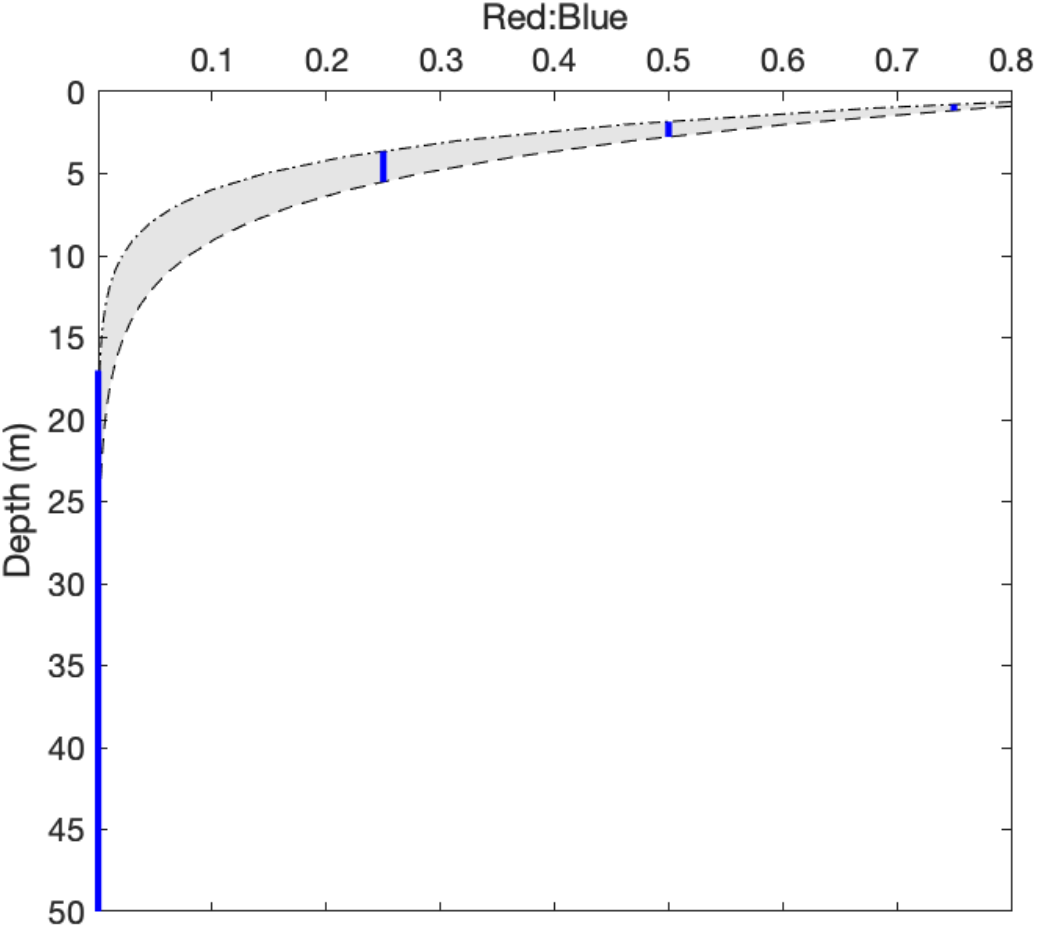
Vertical distribution of the red:blue light ratio in the sea. The grey-shaded region indicates the range of variation for different Jerlov water types (values from Table V of Austin & Petzold, 1986). The blue lines indicate the ratio used in the laboratory experiments shown in Figure 2.

**Figure S4.**
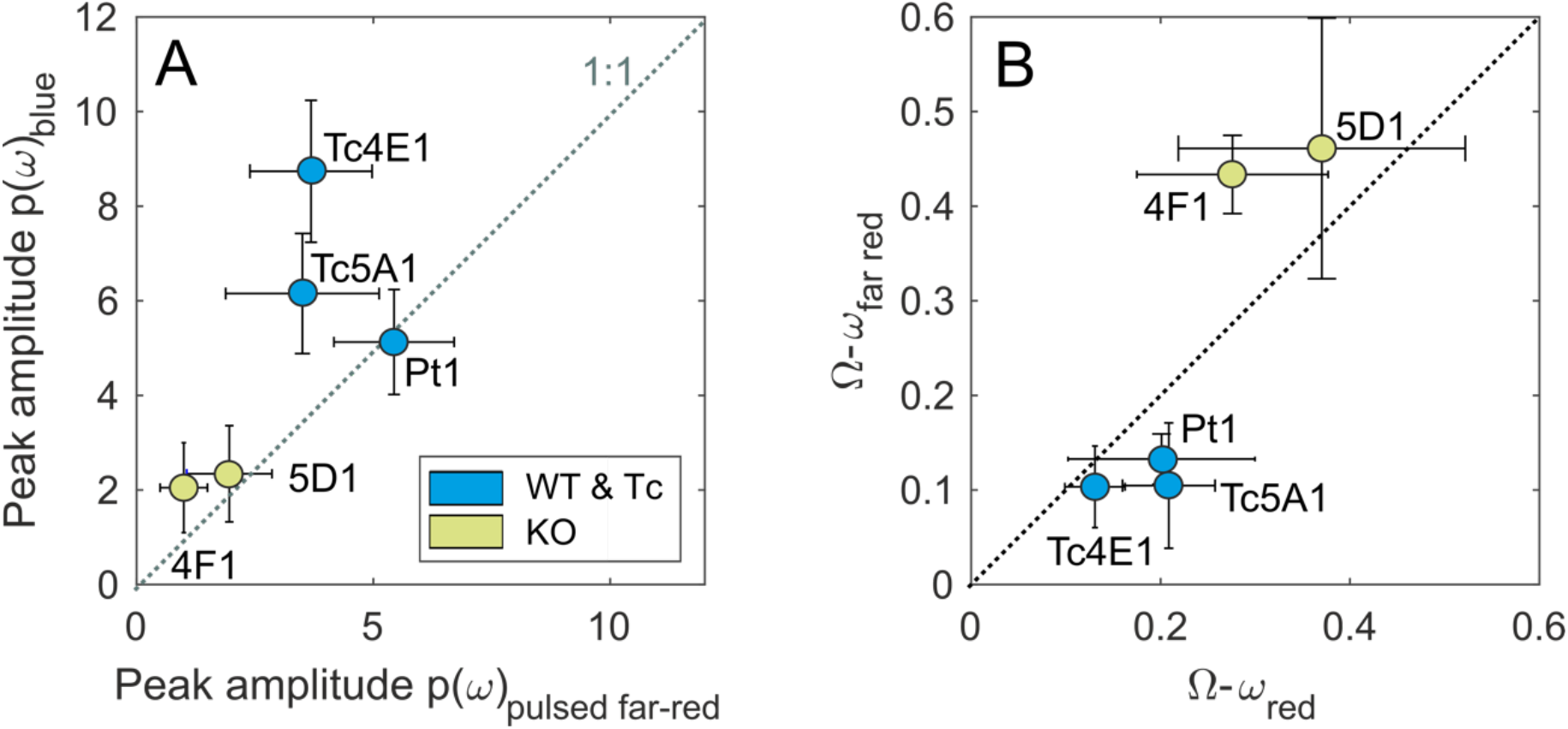
A) Peak amplitude of the power spectrum for experiments with constant blue light compared to pulsed far-red light. B) Compared the emergent frequency difference between the wobble frequency (Ω) and pulsed light-emitting diode (LED) light frequency (ω) for red and far-red experiments.

